# Whole genome CRISPR knockout screen reveals ID3 as a key regulator of myeloma cell survival via TCF3 and c-MYC

**DOI:** 10.1101/2025.09.09.675036

**Authors:** Clara Andersson-Rusch, Md Abu Hanif, Ingrid Quist-Løkken, Pål Sætrom, Miłosz Roliński, Johanna Fredrikke Nordstrand Møen, Naiara Campos Roman, Kristine Misund, Vidar Beisvåg, Per Arne Aas, Barbara van Loon, Toril Holien

## Abstract

Bone morphogenetic proteins (BMP) induce apoptosis in myeloma cells and the mechanism behind this could point to new therapeutic targets. Here, we did a whole genome CRISPR/Cas9 knockout screen using the INA-6 myeloma cell line. Apoptosis was induced with BMP9 and the relative amounts of sgRNAs in treated versus control cells were determined with next-generation sequencing. We identified key positive control genes and a substantial number of novel genes that could be involved in BMP-induced apoptosis. One of the overrepresented genes was the known BMP target gene *ID3*. We found that ID3 was potently induced by BMP9 treatment and that depletion of ID3 protected cells from c-MYC downregulation and apoptosis. ID3 is known to heterodimerize with basic helix-loop-helix (bHLH) TCF transcription factors. In the screen, *TCF3*, *TCF4*, and *TCF12* were among genes that potentially protected cells from apoptosis. Knockdown of *TCF3*, and to some extent *TCF12*, led to lower basal c-MYC levels and lower cell viability, and this was more pronounced after BMP9-treatment. Our results suggest that ID3 plays an important role in regulating the survival of myeloma cells, at least in part by forming heterodimers with TCF3 and thus preventing expression of the c-MYC oncogene.

## Introduction

Multiple myeloma is an incurable cancer of the plasma cells that arises from plasma cells. The disease is heterogeneous and is characterized by hypercalcemia, renal failure, anemia and lytic bone disease^1^. Efficient treatment options exist for multiple myeloma and most patients initially respond very well. Nevertheless, patients become resistant over time and relapse, and the remissions last shorter for each new treatment regimen^1^. Many of the commonly used drugs aim to induce programmed cell death or apoptosis in the myeloma cells^2^. Detailed knowledge about mechanisms that regulate myeloma cell apoptosis could promote the development of novel and more efficient drugs.

Bone morphogenetic proteins (BMPs) are a subgroup of ligands in the transforming growth factor (TGF)-β family. BMP signaling is initiated when ligands form heterotetrameric complexes with two type I and two type II receptors, leading to activation of the SMAD1/5/8 pathway. The activation of SMAD1/5/8 by phosphorylation leads to the formation of a complex consisting of two SMAD1/5/8 molecules and one SMAD4 molecule^3^. This complex translocates into the nucleus and cooperates with other transcription factors to regulate the expression of target genes. Although active SMAD complexes bind weakly to DNA on their own, they are involved in the regulation of several genes^4^. Among the most recognized BMP-SMAD1/5/8 target genes are the genes that encode the inhibitors of differentiation (ID)1-4 helix-loop-helix (HLH) proteins^5^. The expression of the ID genes is quickly and potently induced by SMAD1/5/8 activation and therefore used in reporter assays to detect BMP activity^6, 7^.

BMPs induce growth arrest and apoptosis in myeloma cells both *in vitro* and *in vivo* via the BMP type I receptors ALK2 (*ACVR1*) and ALK3 (*BMPR1A*)^8–10^. Multiple myeloma cells are dependent on the c-MYC oncogene for survival,^11^ and we have shown that BMPs induce apoptosis in myeloma cells through SMAD-dependent downregulation of c-MYC^12^. However, the detailed mechanism between SMAD activation and c-MYC downregulation is not known.

CRISPR pooled libraries are powerful tools for unbiased discovery and functional characterization of specific genetic elements. Positive screens are widely used to enrich and gather resistant cells and can be used to identify genes related to the mechanisms that drive apoptosis^13^. Likewise, negative screens can be used to identify genes that protect cells from apoptosis. Such functional screens have successfully identified drug resistance mechanisms in hematological cancers, including multiple myeloma^14^. We here used a whole-genome lentiviral pooled CRISPR/Cas9 knockout (KO) screen to identify genes affecting apoptosis in the multiple myeloma cell line INA-6 treated with BMP9. The screen resulted in several interesting gene hits, and out of these we focused on the potential role of the known SMAD1/5/8-target gene *ID3* in BMP-induced apoptosis.

The basic helix-loop-helix (bHLH) proteins are part of a family of transcription factors with essential roles in several processes, including development of the nervous system and muscles^15^. Members of the bHLH family rely on homo- or heterodimerization for transcriptional activity by binding to ‘E-box’ (CANNTG) sequences in DNA. ID3, and the other ID family proteins, belong to class V of HLH proteins, which lack the basic element responsible for DNA-binding^15^. ID family proteins heterodimerize with other bHLH proteins, inhibiting their DNA-binding ability^16–18^. We found that sgRNAs for genes encoding TCF transcription factors (TCF3, TCF4, and TCF12) were underrepresented in the screen, meaning that they potentially protect the cells from BMP-induced apoptosis. Interestingly, the transcriptional function of the TCF proteins is known to be inhibited by ID proteins^19–21^. Using siRNA, we show that TCF3, and to a lesser extent, TCF12, protects cells from BMP9-induced apoptosis. We propose that increased levels of ID3 inhibits TCFs, leading to lower MYC levels and apoptosis. Our results indicate that ID3 plays a central role in the regulation of myeloma cell survival and that there is a potential to exploit the function of ID3 in the development of novel therapies.

## Materials and methods

### Cells and reagents

The human myeloma cell lines used were INA-6 (a kind gift from Dr Martin Gramatzki, University of Erlangen-Nurnberg, Erlangen, Germany)^22^, IH-1^23^ and KJON^24^ (both established in our lab), RPMI-8226 (ATCC, Rockville, MD, USA) and Karpas-417 (ECACC, Sigma-Aldrich, Oslo, Norway). We also used the diffuse large B-cell lymphoma (DLBCL) cell line DOHH-2 (Leibniz Institute DSMZ, Braunschweig, Germany). Recombinant human BMP9 (#3209-BP) and BMP10 (#2926-BP) were from R&D Systems (Bio-Techne, Abingdon, UK).

### Lentiviral library generation

The generation of lentiviral library was performed as shown before and a detailed description is provided in Supplementary Materials and methods^25^.

### Genome-wide CRISPR-Cas9 screen

287 × 10^6 INA-6 cells were seeded at 1.4 × 10^6 cells/mL in growth medium in the presence of 8 µg/mL polybrene (#107689, Sigma-Aldrich), and lentivirus GeCKOv2 library A and B was added at a multiplicity of infection (MOI) of 0.25 and incubated at 37 ℃ for 4 h before the cells were diluted to 0,35 × 10^6 cells/mL. The number of transduced cells was calculated to achieve at least 700 × coverage per single guide (sg) RNA construct. Puromycin (0.5 µg/mL) was added after 48 h and the selection for transduced cells was continued for 5 days. Cells were then expanded and 5 × 10^7 cells in triplicates were collected as pre-treatment control. The remaining cells were divided into three replicates for each treatment arm and treated for 3 or 7 days with or without BMP9 (0.2 ng/mL). For each condition up to 5 × 10^7 cells were collected using Optiprep (Axis-Shield, Oslo, Norway) for density gradient centrifugation to remove dead cells. Genomic DNA was isolated using DNA Isolation Kit for Cells and Tissues (11814770001, Roche). Illumina sequencing libraries were prepared using a two-step PCR method with Herculase II fusion DNA polymerase (#600679, Agilent). The primers are listed in Supplementary Table S1. For the first PCR, 132.5 µg gDNA was amplified. Each reaction contained 2.5 µg gDNA in a final volume of 100 µL and was cycled as follows: 1 cycle 120 s at 95 ℃; 25 cycles 20 s at 95 ℃; 20 s at 67 ℃; 30 s at 72 ℃; and 1 cycle 180 s at 72 ℃. The second PCR contained 5 µL pooled amplicons from the first PCR in a final volume of 100 µL. The samples were cycled as follows: 1 cycle 120 s at 95 ℃; 10 cycles 20 s at 95 ℃; 20 s at 67 ℃; 30 s at 72 ℃; and 1 cycle 180 s at 72 ℃. For each sample 11 reactions were prepared. Samples were purified using Agencourt AMPure XP beads (A63880, Beckman) and analyzed on the Caliper LabChip GX Nucleic Acid Analyzer (PerkinElmer). Sequencing libraries were denatured, diluted and pooled according to the standard Illumina protocols. Pooled libraries (2.7 nM) were sequenced on a NextSeq 500 high-output flow cell at 81 cycles for the insert and 9 cycles for the index). PhiX DNA (20 %) was spiked into the libraries to increase diversity. FASTQ files were created with bcl2fastq 2.20.0.422 (Illumina, CA, USA).

### Bioinformatic analysis

The software cutadapt (version 1.15) was used to remove the GeCKO 5’ and 3’ flanking sequences from the reads in the FASTQ files and bowtie2 (version 2.3.4.1) was used to align the resulting sequences against the GeCKO FASTA gRNA file. The resulting number of reads aligning per gRNA sequence per library was then used as input for statistical analysis in the voom-limma framework. Specifically, the matrix of counts per gRNA per library was normalized to reads per million (RPM) and analyzed by linear modelling to identify individual gRNAs where the number of reads differed significantly between either BMP9 and control at day 3 or day 7.

The results from the individual gRNA analyses were combined into gene-level compound results by using the following three complementary methods: 1) Stouffer’s method of combining multiple p-values, with the null-hypothesis being that none of the gRNAs targeting a gene are significant; 2) a Student’s *t*-test of whether the mean logFC for all gRNAs targeting a gene is different from the mean logFC for the GeCKO library’s control gRNAs; and, 3) and the result from the robust rank aggregation algorithm test^26^ of whether the gRNAs a gene are enriched towards the top of the ordered list of all gRNAs, with the null-hypothesis being that the gRNAs are randomly ordered.

### Other methods

Detailed descriptions of the remaining methods are provided in Supplementary Materials and methods.

## Results

### CRISPR/Cas9 KO screen revealed *ID3* as a top potential gene necessary for BMP-induced apoptosis

The exact mechanisms leading to BMP-induced apoptosis in multiple myeloma cells are unknown. We therefore performed an unbiased CRISPR/Cas9 based KO screen targeting the entire human genome in the INA-6 myeloma cell line. Many BMPs induce apoptosis in myeloma cells and we chose to use BMP9, one of few BMPs found in physiological concentrations in circulation^27^, and a potent inducer of apoptosis in these cells^10^. The screen is presented schematically (Figure 1A). This type of screen can identify both negative and positive regulators of apoptosis, but we focused on positive hits, meaning genes that are needed for the cells to go into apoptosis. Based on our previous studies, we used a BMP9 dose (0.2 ng/mL) needed to achieve approximately 90 % cell death in control cells (IC90).

**Figure 1.**
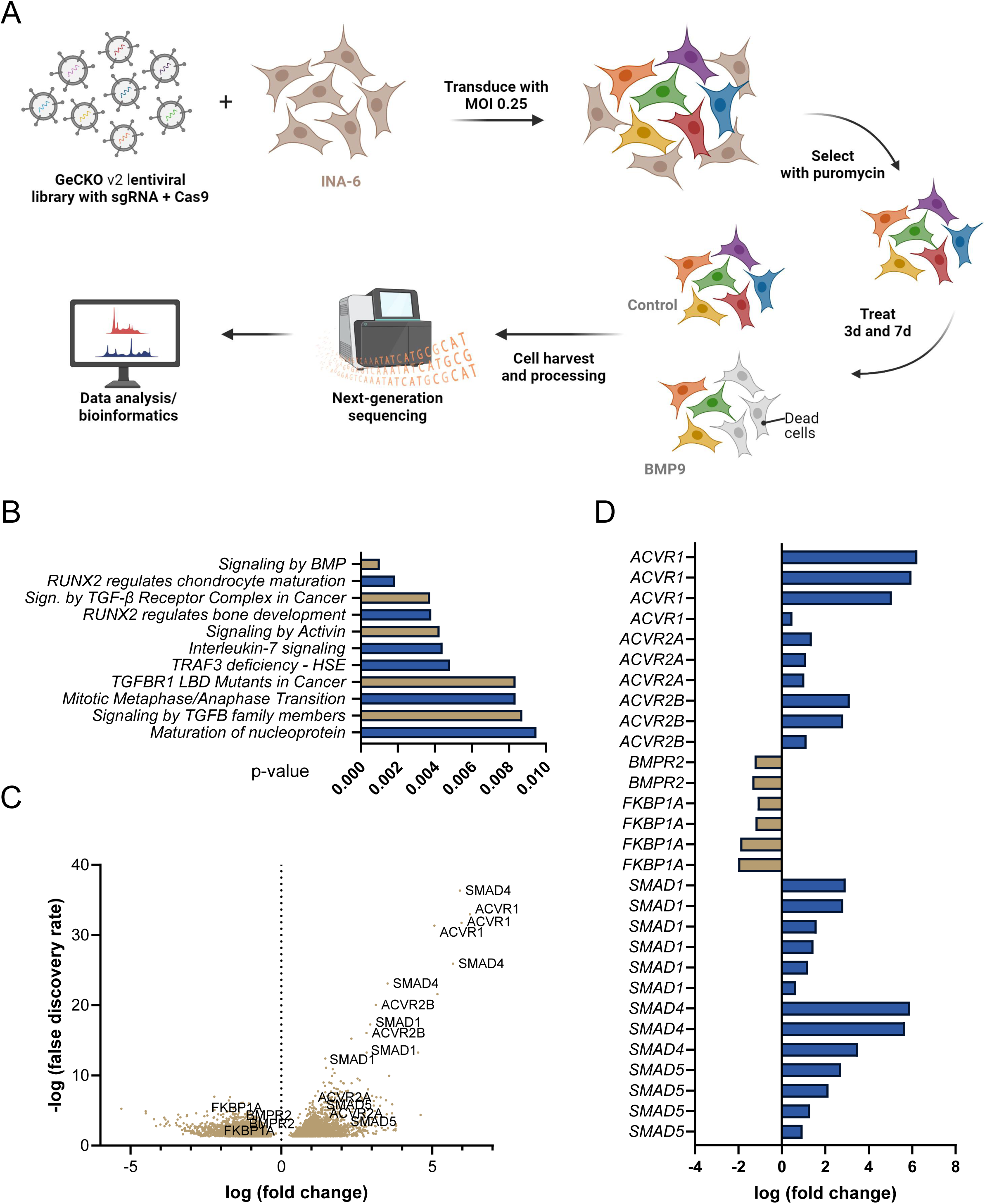
Whole-genome CRISPR/Cas9 knockout screen identified novel genes that either promote or inhibit BMP-induced apoptosis. A. Schematic overview of the CRISPR/Cas9 screen. In short, INA-6 cells were transduced with lentivirus from the GeCKOv2 library with 6 sgRNAs per gene. Transduced cells were selected with puromycin treatment, split into flasks, and treated with or without BMP9 (0.2 ng/mL) in triplicates. Cells were collected at day 3 and 7 of treatment and to avoid background in the NGS analysis, dead cells were removed by density gradient centrifugation. Genomic DNA from the surviving cell population was sequenced using a high-throughput platform, leading to identification of candidate genes. Created with BioRender.com. B. Top Reactome pathways correlating with the 500 most significant sgRNAs found in the BMP9 treated cells versus control on day 7. Gene sets related to BMP signaling are shown in light brown. C. Volcano plot showing enriched single sgRNA that were overrepresented (right) or underrepresented (left) in the BMP9 treated cells on day 7 of treatment. D. The significantly different sgRNAs for positive controls sorted alphabetically. The log fold change numbers are shown and visualized with blue bars showing increased levels compared with control and light brown bars showing decreased levels compared with control.

The efficacy of the chosen dose was confirmed in cell culture flasks to mimic the screen conditions (data not shown). The cells were transduced with the lentiviral library that contained 6 different sgRNAs per gene. To account for the complexity and the high number of chromosomes in the INA-6 cell line^28^, we used a calculated coverage of 700 and multiplicity of infection (MOI) of 0.25. The cells were treated with or without BMP9 in triplicates. Genomic DNA isolated from samples were collected on day 3 and 7 of treatment and processed for next-generation sequencing (NGS) and bioinformatic analysis as described in Materials and methods.

Very few significant differences could be seen on day 3, but on day 7 of treatment > 9000 significantly over- or underrepresented sgRNAs were found in BMP9-treated cells compared with control (Supplementary Table S2). We first used Reactome pathway analysis to map the 500 most significant sgRNAs (Figure 1B). Not surprisingly, “Signaling by BMP” appeared as the most significant pathway involved. Then, all sgRNAs were plotted as a volcano plot to visualize how single sgRNAs related to each other (Figure 1C). We identified several positive controls, e.g. genes known to be important for BMP-induced apoptosis (*ACVR1*, *ACVR2B*, *SMAD1*, and *SMAD4*) and genes that inhibit SMAD1/5-activity in these cells (*BMPR2* and *FKBP1A*)^10, 12, 29–32^. The log2 fold changes for these sgRNAs are shown in Figure 1D. The identification of these sgRNAs confirmed that the screen had worked as planned and that the unknown genes/sgRNAs in our list were of potential relevance. We also analyzed the data using a compound approach, looking at all 6 sgRNAs per gene together, and how each gene was differently represented in BMP9-treated versus control cells. This approach resulted in 62 significant (p < 0.05) gene hits (Supplementary Table S3). Among the top genes from the compound analysis, excluding positive controls, *ID3* stood out as an interesting hit with unresolved function in myeloma cell apoptosis.

### BMP9 induced the expression of ID3 in different cell lines

ID3 is part of the family of HLH proteins that are named ID1-4. In myeloma cells, *ID2* is the one with the highest baseline mRNA expression, whereas *ID1* and *ID3* are expressed, but at low levels, and *ID4* is basically not expressed (Figure 2A) (data from Keats Laboratory Database, https://www.keatslab.org). *ID3*, but also *ID1*, are recognized as SMAD1/5 target genes^33^. Therefore, we used an *ID1* reporter cell line, INA-6 BRE-luc, that we generated for rapid measurement of BMP-SMAD activity, and observed that BMP9 potently induced luciferase expression^34^ (Figure 2B). We then wanted to compare reduced cell viability and induction of *ID3* and *ID1* after BMP9 treatment in a broader set of cell lines. The myeloma cell lines INA-6, IH-1, Karpas-417, KJON, and RPMI-8226, as well as the BMP-sensitive DLBCL cell line DOHH-2 were treated with BMP9 for 72 h and relative cell viability was measured (Figure 2C). Out of these, RPMI-8226 was the least sensitive of these cell lines to BMP9-induced apoptosis. We then measured mRNA expression of *ID3* and *ID1* in the same cell lines after 4 hours of BMP9 treatment (Figure 2D, E). For comparison, *ID3* and *ID1* levels were also measured in cells treated with a different BMP, BMP10 which we have previously shown do not induce SMAD signaling in INA-6 cells (Supplementary Fig. S1)^32^. There was a clear increase in the expression of both *ID3* and *ID1* in all cell lines tested, except for RPMI-8226, which is consistent with that cell line not being very BMP9-sensitive. We then compared SMAD1/5 phosphorylation in two cell lines where BMP9-treatment led to reduced cell viability and strong induction of *ID3* and *ID1* (KJON-1 and INA-6), with RPMI-8226, where BMP9-treatment had limited effect. Although BMP9-treatment led to phosphorylation of SMAD1/5 in RPMI-8226, the levels were lower compared with KJON-1 and INA-6 (Figure 2F, and 2G for quantification). The results are consistent with our previous data showing that RPMI-8226 cells express less of the BMP9-receptor *ACVR1* than KJON-1 and INA-6^30^. In contrast, BMP10 only activated SMAD1/5 in KJON cells, consistent with KJON-1 being the only cell line out of these three that expresses *BMPR1A*, which is a type I receptor for BMP10^30^. Our results suggest a relation between SMAD-activation, induced expression of ID3 (and ID1), and reduced cell viability, and that variations in sensitivity could at least partly be explained by the expression of different BMP receptors in the cells tested.

**Figure 2.**
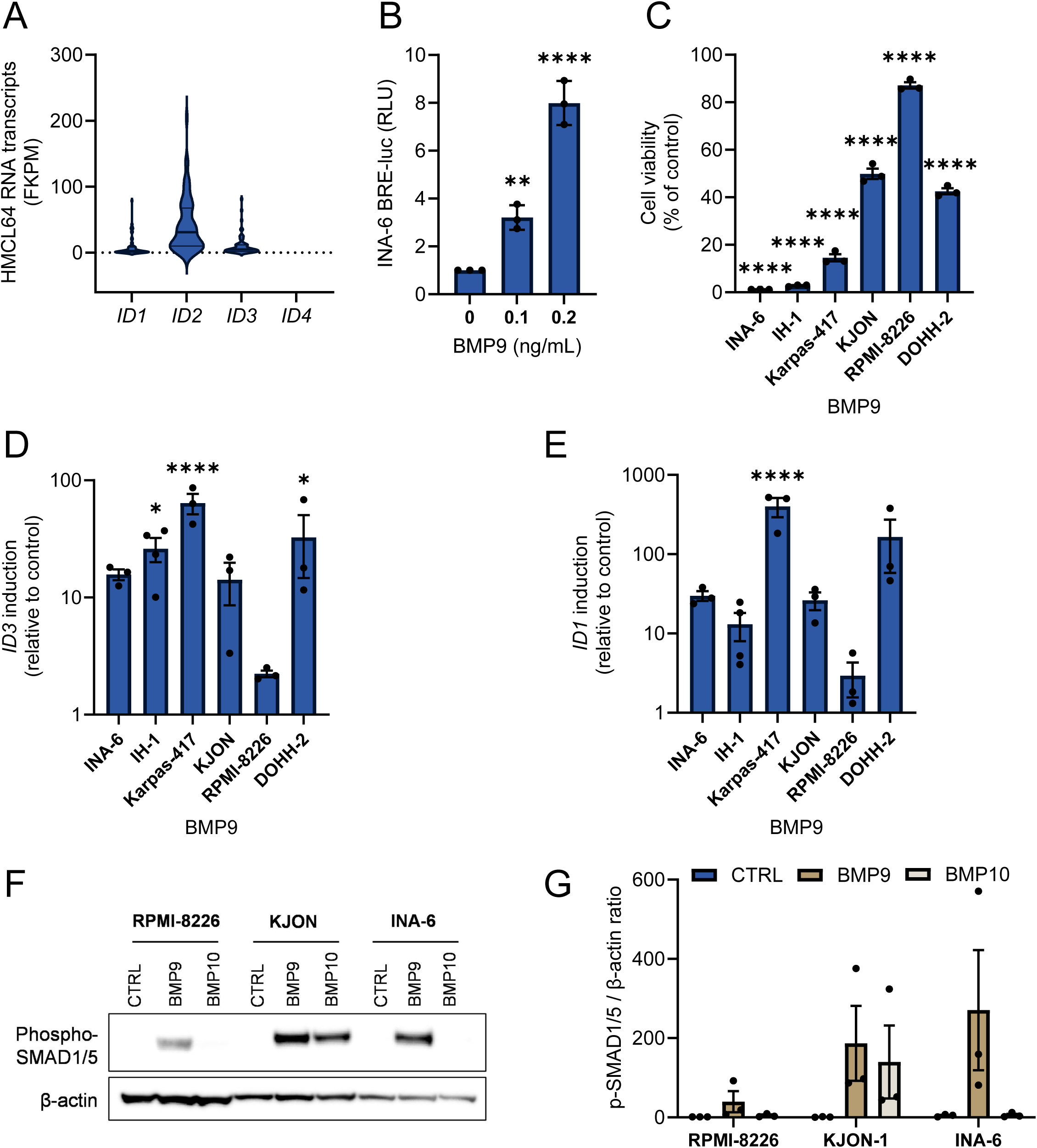
BMP9 induced the expression of *ID3* in different cell lines. A. The transcript levels of ID1-ID4 in 64 human myeloma cell lines (HMCL) were obtained from Keats Laboratory database (https://www.keatslab.org) and are shown as fragments per kilobase per million (FKPM) using a violin plot. B. INA-6 BRE-luc cells were treated for 18 h with BMP9 (0, 0.1 and 0.2 ng/mL) before luciferase activity was measured using Britelite plus reporter assay. The mean and SEM of n=3 independent experiments were plotted (**; p<0.01, ****; p<0.0001, one-way ANOVA, Dunnett’s multiple comparisons test). C. Different cell lines were treated with BMP9 (50 ng/mL) for 72 h before relative cell viability was measured using CellTiter Glo. Cell viabilities were plotted relative to medium control. The same cell lines as in C were treated with BMP9 (20 ng/mL) for 1 h and the mRNA levels of *ID3* (D) and *ID1* (E) were determined by RT-qPCR and the comparative Ct method with GAPDH as housekeeping gene. All graphs C-E represent mean and SEM of n=3 independent experiments, *; p<0.05, ****; p<0.0001, two-way ANOVA, Šídák’s multiple comparisons test. E. RPMI-8226, KJON-1, and INA-6 cells were treated with BMP9 (20 ng/mL) or BMP10 (50 ng/mL) for 1 h before measuring phospho-SMAD1/5 activation using western blot, with β-actin as loading control. Shown is a representative of n=3 independent experiments. F. Combined phospho-SMAD1/5 /β-actin ratios of all three independent the western blots from E.

### Loss of ID3 counteracted BMP-induced apoptosis and c-MYC downregulation

To further validate and investigate the role of ID3 in BMP-induced apoptosis, we generated two independent INA-6 ID3 KO cell clones using CRISPR/Cas9, each with different sgRNAs targeting ID3, KO1 and KO2, and non-targeting control (NTC) cells. Normally, ID3 protein is barely detectable in these cells in culture but increases rapidly when SMAD1/5/8 is activated. Loss of ID3 expression was therefore shown by western blot of cells treated with BMP9 (5 ng/mL) for 4 h (Figure 3A, and Supplementary Fig. S2A for quantification). In wildtype (WT) and NTC cells there was a clear BMP9-induced ID3 protein band, whereas in the ID3 KO clones we could not detect ID3. Of note, a smaller band was detected in the ID3 KO2 cells, suggesting that a truncated variant of ID3 was expressed in this clone. The cells were then treated with increasing doses of BMP9 for 72 h, and apoptosis was analyzed using annexin V/PI and flow cytometry (Figure 3B). We also analyzed cell viability with the sensitive CellTiter Glo ATP assay (Figure 3C). The ID3 KO clones were less sensitive to BMP9 than WT and NTC cells, and the ID3 KO1 clone was the least sensitive of the two and was used for further analyses.

**Figure 3.**
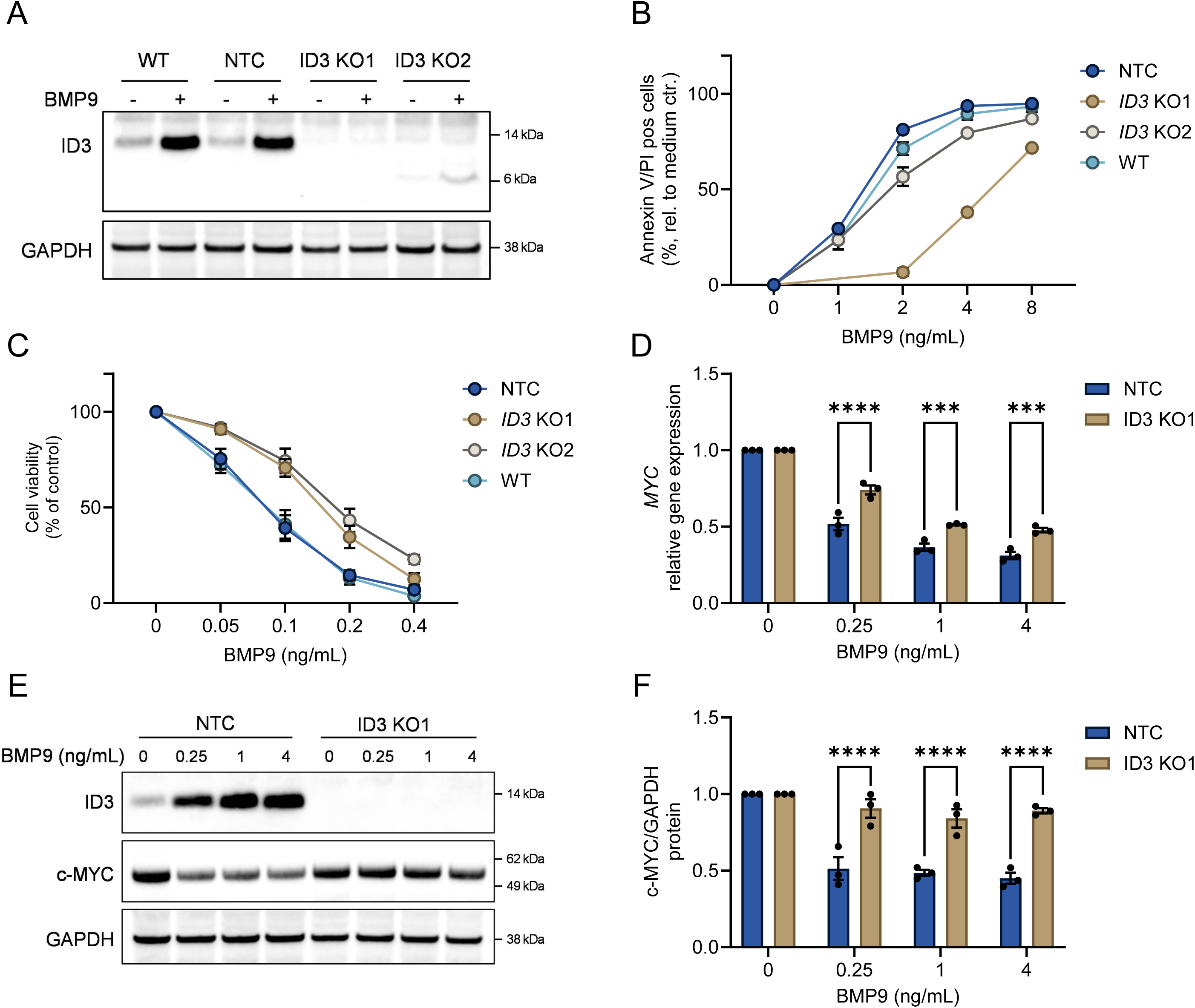
Loss of ID3 counteracted BMP-induced apoptosis and c-MYC downregulation. A. INA-6 WT, NTC, ID3 KO1, and ID3 KO2 were treated with BMP9 (5 ng/mL) for 4 h before analysis of ID3 protein levels by western blot with GAPDH as a loading control. B. INA-6 variants were treated with BMP9 for 72 h and the relative cell viabilities were determined with CellTiter Glo. C. INA-6 variants were treated with BMP9 for 72 h. Cells were stained with annexin V/FITC and propidium iodide (PI) and positive cells relative to medium control were plotted. The graphs were plotted relative to medium control and each bar represents the mean and SEM of n=3 independent experiments. D. MYC gene expression in INA-6 NTC and ID3 KO1 cells was quantified with RT-qPCR after treatment with different doses of BMP9 for 4 h. The graph shows the mean and SEM of n=3 independent experiments. The comparative Ct method was used and GAPDH was used as housekeeping gene. E. INA-6 NTC and ID3 KO1 cells were treated with different doses of BMP9 for 6 h. Western Blot was performed to detect MYC and ID3 protein levels and GAPDH as a loading control. Shown is a representative of n=3 independent experiments, and densitometric analysis is shown in F. Densitometric analyses of ID3 levels from the blots in Fig. 3A and 3F are shown in Supplementary Fig. S2. Two-way ANOVA and Sidak’s multiple comparison test was used in D and F (***; p<0.001, ****; p<0.0001).

The *MYC* oncogene is necessary for survival of myeloma cells and SMAD-dependent MYC downregulation leads to apoptosis^11, 12^. We therefore wanted to clarify if ID3 was involved in c-MYC downregulation. INA-6 NTC and ID3 KO1 cells were treated with different doses of BMP9 for 4 h and the mRNA levels of *MYC* were measured with qRT-PCR (Figure 3D), whereas the protein levels of ID3 and c-MYC were determined by western blot after 6 h BMP9 treatment (Figure 3E, and 3F (c-MYC) and Supplementary Fig. S2B (ID3) for quantification). In NTC cells, there was a dose-dependent induction of ID3 protein and a downregulation of *MYC* mRNA and protein, whereas in the ID3 KO1 cells we did not detect ID3, the *MYC* mRNA was significantly less downregulated, and we only saw a minor reduction in c-MYC protein levels. We conclude that ID3 is needed for BMP9-induced c-MYC downregulation and apoptosis in myeloma cells.

### TCF family proteins interact with ID3, and gene expression levels of TCF3 predict overall survival in myeloma patients

We then asked ourselves how ID3 could affect MYC mRNA and protein levels. ID proteins are HLH transcription factors, a big group of proteins that often interactions to regulate gene transcription. We therefore made an overview of the HLH proteins that were significantly over- or underrepresented in the screen (Supplementary Fig. S3). We also mapped known physical interactors of ID3 using the STRING database (Figure 4A). Only six proteins were identified: PAX5, PUF60, SREBF1, TCF3, TCF4, and TCF12. All three TCFs were represented with 1-3 sgRNAs with a negative fold change (Supplementary Fig. S3 and Table S2), indicating that they may protect the cells from BMP-induced apoptosis. In contrast, PUF60 showed up with one sgRNA with positive fold change, and PAX5 and SREBF1 were not detected in the screen (Supplementary Table S2), and we decided to focus on the TCFs. The three TCF proteins are bHLH proteins, also known as the E protein transcription factor family, or class I HLH transcription factors^15^. ID family proteins are known to antagonize the class I bHLH proteins via physical interaction that prevents DNA binding. All TCFs are expressed in myeloma cells, with TCF3 having the strongest expression (Figure 4B).

**Figure 4.**
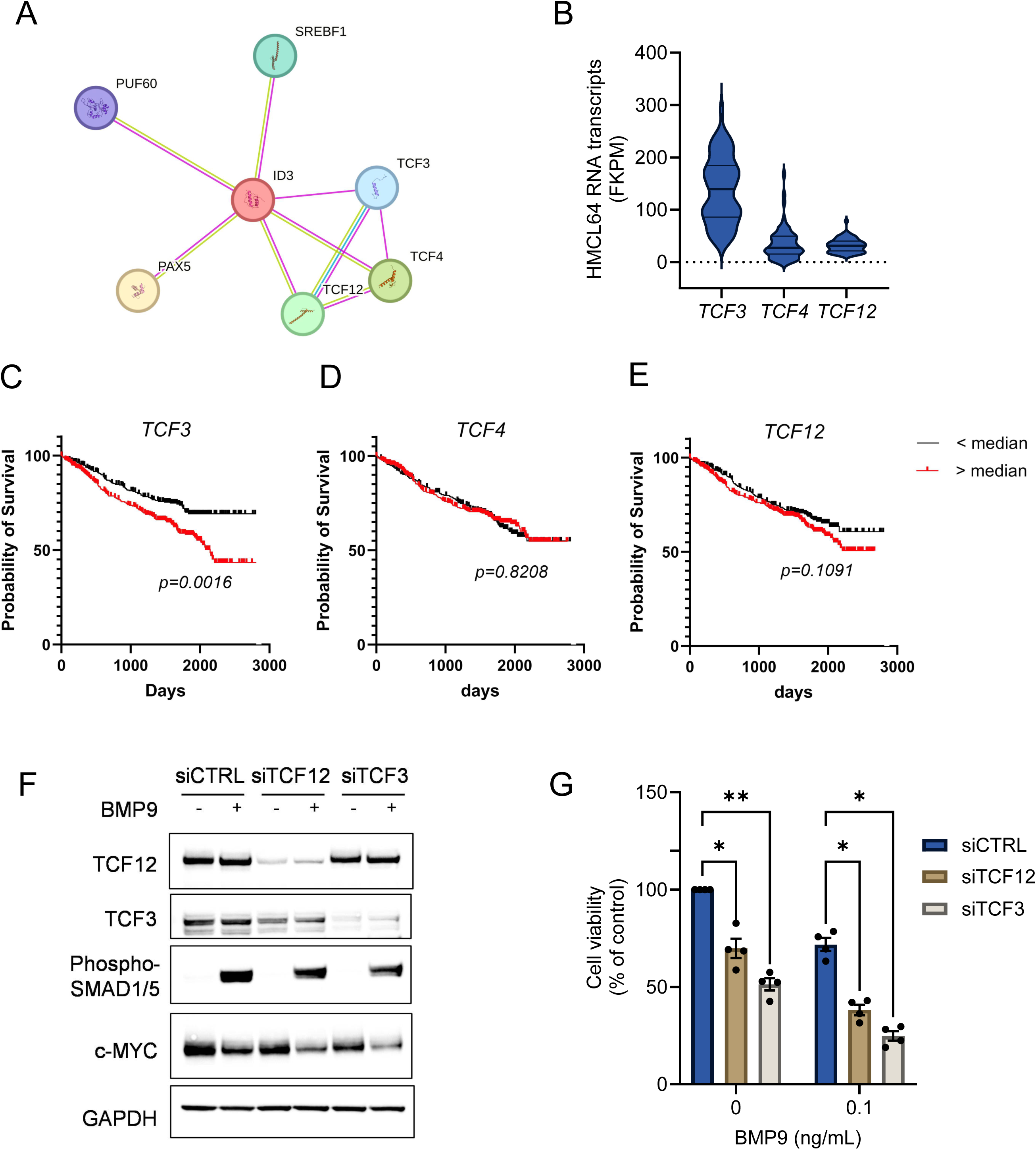
TCF family proteins interact with ID3, and TCF3 transcript levels predict overall survival in myeloma patients. A. Known protein-protein interactions of ID3 from the STRING database showing only known physical interactions. B. The transcript levels of TCF3, TCF4, and TCF12 in 64 human myeloma cell lines (HMCL) were obtained from Keats Laboratory database (https://www.keatslab.org) and are shown as fragments per kilobase per million (FKPM) using a violin plot. C-E. Overall survival curves generated from the MMRF CoMMpass data (IA16 release, n=767) by comparing gene expression above and below median for TCF3, TCF4 and TCF12. The results were generated from the MMRF-CoMMpass database, n = 767, high>upper quartile, low<lower quartile. The Log-rank Mantel-Cox test was used to test for significance. G. INA-6 cells were treated with siRNA targeting TCF3, TCF12 or with non-targeting control siRNA. After treatment for 2 h with BMP9 (1 ng/mL), the protein levels of TCF12, TCF3, phospho-SMAD1/5/8, and c-MYC were determined and compared with GAPDH as a loading control. F. The same siRNA-treated cells as in F were incubated with BMP9 for 48 h and relative viability was measured with CellTiter Glo. The graph shows mean and SEM of n=3 independent experiments. Two-way ANOVA with Tukey’s multiple comparisons test was used to test significance (*; p<0.05, **; p<0.01, ***; p<0.001).

Moreover, when we used DepMap portal to compare selected genes and cancers, TCF3 stood out as a gene that myeloma cells depend on for survival, and the dependency is relatively specific for lymphoid tissue, in contrast to MYC that is a general cancer dependency gene (Supplementary Fig. S4). We used the MMRF-CoMMpass data set to see if there was any correlation between overall survival and each of the genes *TCF3*, *TCF4* and *TCF12* (Figure 4C-E). We found that low *TCF3* expression correlated with prolonged overall survival, whereas TCF4 and TCF12 showed no correlation with overall survival. Knockdown of TCF3 and TCF12 using siRNA supported a role for TCF3, and to a lesser extent TCF12, in regulation of c-MYC levels (Figure 4F) and cell survival (Figure 4G) in the INA-6 myeloma cell line. Taken together, we here show that ID3 is a key regulator of c-MYC levels and as well as myeloma cell survival downstream of BMP-SMAD activation, possibly by binding and inhibiting the TCF3 transcription factor (illustrated in Figure 5). TCF3 regulates c-MYC levels and low levels of TCF3 predicts prolonged patient survival.

**Figure 5.**
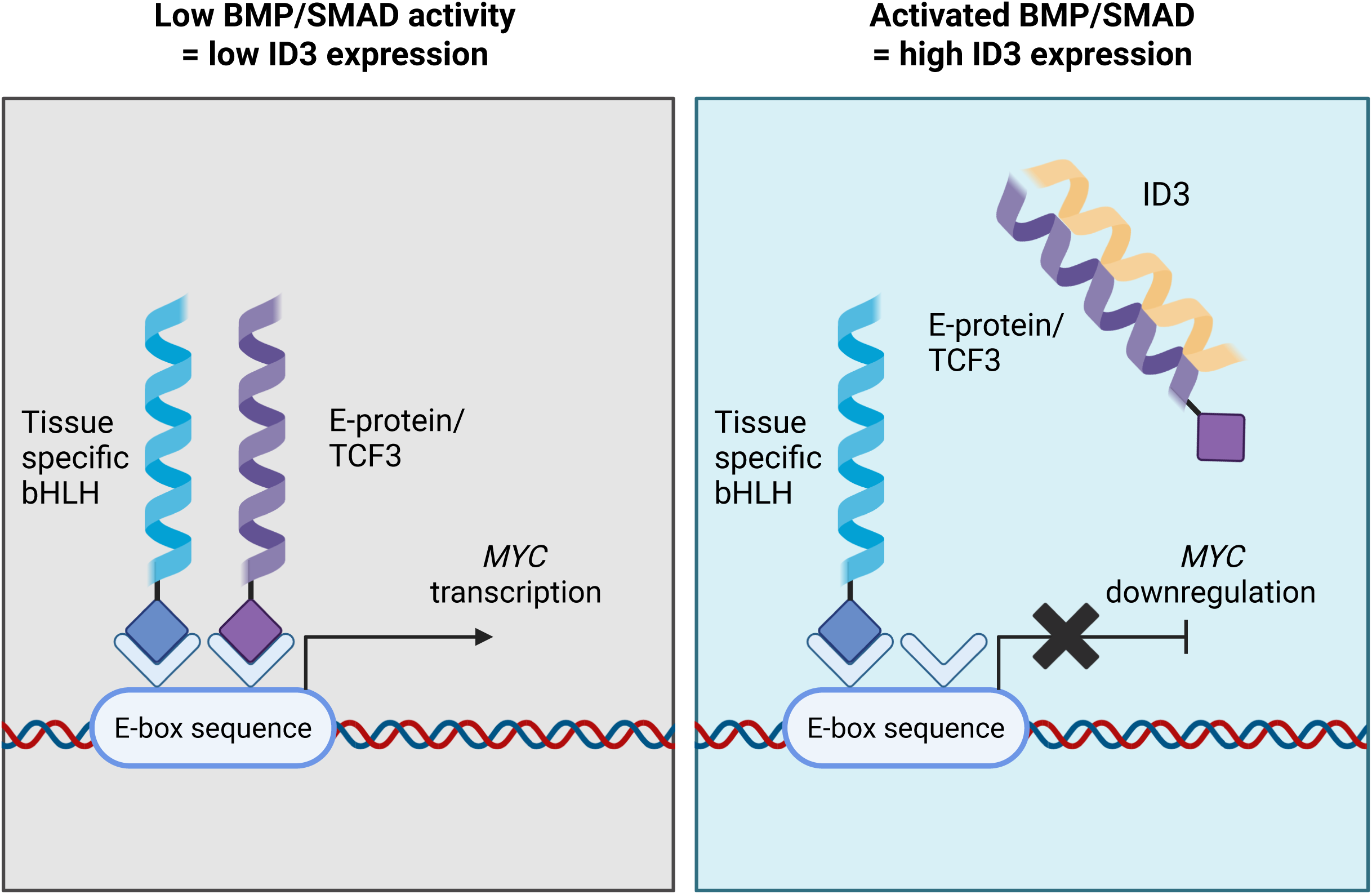
Proposed mechanism of BMP/SMAD-induced downregulation of *MYC* and induction of apoptosis in myeloma cells. In the absence of BMP/SMAD1/5/8 activity (BMP/SMAD) E-proteins such as TCF3, heterodimerize with tissue specific bHLH proteins (alternatively homodimerize) and positively regulate transcription of *MYC*. Activated BMP/SMAD signaling leads to a quick and potent upregulation of ID3, a HLH protein lacking the DNA-binding domain. ID3 then heterodimerizes with the TCF3 transcription factor and prevents its binding to DNA, resulting in downregulation of *MYC* and myeloma cell apoptosis. The figure was made with Biorender.com.

## Discussion

BMPs induce apoptosis in multiple myeloma cells and knowledge about this mechanism could be useful in development of novel drugs that target myeloma cells. In this study we used a CRISPR/Cas9-based whole genome screen and identified ID3 as an important mediator of BMP-induced apoptosis. The inhibitors of differentiation (ID)-proteins, ID1-4, are a family of helix-loop-helix (HLH) proteins that are known SMAD target genes^6, 7, 35^. Although these proteins have many similarities, their functions are not completely overlapping, and they have different roles depending on the cellular context. Earlier studies have found that ID3 can have an important role in regulating cell proliferation in B-lymphocytes^36^. For instance, ID3 inhibited B cell development *in vitro*^37^ and was required for growth inhibition mediated by TGF-β in pro-B cells ^36^. Using exome sequencing, ID3 was found to be mutated in 34 % of Burkitt lymphoma cases but not in DLBCL^38^. Later studies have found recurrent alterations of ID3 in high-grade B-cell lymphoma and marginal zone lymphomas with histologic transformation^39, 40^. Taken together, many studies suggest a potential role of ID3 as a tumor suppressor gene in hematological cancers.

ID3 and ID1 are often poorly expressed in multiple myeloma cells at baseline. These genes are however quickly induced upon stimuli, such as via activated SMAD1/5-SMAD4 transcription complexes that bind to BMP-responsive elements in the ID gene promoters^6, 7,35^. In the cell lines tested here, we found that induction of ID3 correlated well with reduced cell viability mediated by BMP treatment. The cell line that was least sensitive to BMP9, RPMI-8226, also showed poorer response to BMP9-induced ID-induction. This coincides with the expression of BMP9 receptors, as we earlier measured that RPMI-8226 expresses lower levels of ACVR1, ACVR2A and ACVR2B than cell lines INA-6, IH-1, and KJON-1^30^. This observation is not surprising as the minimum requirement for activation of SMADs is the presence of appropriate receptors on the target cells.

SMAD activity in myeloma cells has previously been associated with MYC downregulation, but the mechanism leading to this has been unclear. One of the most known MYC interactors is its partner protein MAX^41^. Both MYC and MAX are class III/IV bHLH proteins that can heterodimerize with each other, bind E-box sequences and regulate the transcription of a wide range of target genes^41^. When we knocked out ID3, the cells were protected against BMP9-mediated cell death. ID3 KO also protected cells against MYC downregulation on both mRNA and protein level. Heterodimerization of ID3 with MYC could in theory be a potential mechanism for BMP-induced apoptosis. However, ID3 has not been reported to be able to bind either MYC or MAX. This led us to suspect one or more other downstream players involved with MYC downregulation. Several members of the bHLH superfamily, such as certain E-box proteins, hetero- or homodimerize to be able to bind DNA^15^. It is well established that the ID proteins can dimerize with the bHLH transcription factors TCF3, TCF4 and TCF12 and block their DNA binding ability^42^, as we also found using STRING network analysis. TCF3, TCF4 and TCF12 are dimeric transcription factors that bind to E-box sequences and are proposed to play important roles in several cellular processes^15^. The ability to heterodimerize with tissue-specific bHLH proteins is important for the diverse roles of TCFs^42^. In pediatric B-lymphoblastic leukemia and lymphoma, TCF3 rearrangements leading to gene fusions are recurrent, with TCF3 fusion to PBX1 being the most common^43^.

In this setting, the TCF3 fusion products are thought to be oncogenic drivers. Interestingly, TCF3 was one of 23 myeloma-specific essential genes found via analyses of DepMap project data and the expression of TCF3 correlated with progression-free survival^44^. Another study also found TCF3 to correlate with both progression-free and overall survival in myeloma patients in different data sets^45^. From the available CoMMpass data we also found that low TCF3 expression highly correlated with prolonged overall survival, whereas TCF4 and TCF12 levels did not correlate. TCFs have tissue-specific roles and regulate different genes via transcription and although a direct link between expression and survival was not seen for TCF4 and TCF12, they may still play important roles in this context. Interestingly, proteomic analyses of MYC interactors have suggested that TCF3 can bind MYC directly^46^. Moreover, TCF12 was one of a group of transcription factors found to bind cis-regulatory elements of the myeloma survival factors MYC, PRDM1, POU2AF1, SDC1, and PIM2^45^. Based on our findings and other people’s work we suggest that TCF3, and possibly TCF12 and TCF4, might play important roles in multiple myeloma.

Of note, a limitation of our study is that the BMP effects were only assessed *in vitro*. It is likely that microenvironmental factors will impact the BMP effect. In line with this, a previous study using a mouse myeloma cell line in coculture with the HS-5 human bone stromal cell line blunted BMP-6 induced growth inhibition ^47^. Still, using an adeno-associated virus expressing BMP4 in the liver of a xenograft mouse multiple myeloma model, we have previously shown that tumor load was reduced compared with control mice ^9^. Additionally, the whole genome KO screen was performed in only one cell line. This approach increases both the chances of picking irrelevant hits and of missing potentially interesting genes.

The results presented here suggest that BMPs induce apoptosis via upregulation of ID3 which in turn inhibits the DNA-binding ability of TCF3, antagonizing its ability to bind to the E-box sequence in MYC or a MYC promoting region (Figure 5). Although there are still unresolved questions, we here show that ID3 is a key factor that regulates MYC levels in multiple myeloma cells, possibly via inhibition of TCF3, a survival factor in myeloma cells. This finding could have implications for the development of future therapies, although more studies are needed to characterize this mechanism in detail.

## Acknowledgments

The authors are grateful for technical assistance from Hanne Hella and Nur Mahammad. Next-generation sequencing service was provided by the Genomics Core Facility (GCF), Norwegian University of Science and Technology (NTNU). The bioinformatics analyses were performed at the Bioinformatics core facility (BioCore), NTNU. GCF and BioCore are both funded by the Faculty of Medicine and Health Sciences at NTNU, and Central Norway Regional Health Authority. The study was funded by the Liaison Committee for education, research, and innovation in Central Norway.

## Author contributions

Conceptualization and study design: C.A.R., P.A.A., K.M., V.B., B.v.L., T.H. Performing experiments, collecting, or analyzing data: C.A.R., M.A.H., I.Q.L., P.S., M.R., J.F.N.M., N.C.R., P.A.A., T.H. Preparing figures and writing original draft: C.A.R. and T.H. Reviewing and editing the draft: All authors. Supervision and funding acquisition: B.v.L. and T.H. All authors read and approved the final manuscript.

## Data availability statement

All data generated or analyzed during this study are included in this published article and its supplementary information files.

## Notes

### Competing Interest Statement

The authors have declared no competing interest.

## References

1. Rajkumar SV. Multiple myeloma: 2024 update on diagnosis, risk-stratification, and management. Am J Hematol 2024 Sep; 99(9): 1802–1824.

2. Teh CE, Gong JN, Segal D, Tan T, Vandenberg CJ, Fedele PL, et al. Deep profiling of apoptotic pathways with mass cytometry identifies a synergistic drug combination for killing myeloma cells. Cell Death Di1er 2020 Jul; 27(7): 2217–2233.

3. Shi Y, Massagué J. Mechanisms of TGF-beta signaling from cell membrane to the nucleus. Cell 2003 Jun 13; 113(6): 685–700.

4. Morikawa M, Koinuma D, Miyazono K, Heldin CH. Genome-wide mechanisms of Smad binding. Oncogene 2013 Mar 28; 32(13): 1609–1615.

5. Hollnagel A, Oehlmann V, Heymer J, Rüther U, Nordheim A. Id genes are direct targets of bone morphogenetic protein induction in embryonic stem cells. J Biol Chem 1999 Jul 9; 274(28): 19838–19845.

6. Korchynskyi O, ten Dijke P. Identification and functional characterization of distinct critically important bone morphogenetic protein-specific response elements in the Id1 promoter. J Biol Chem 2002 Feb 15; 277(7): 4883–4891.

7. López-Rovira T, Chalaux E, Massagué J, Rosa JL, Ventura F. Direct binding of Smad1 and Smad4 to two distinct motifs mediates bone morphogenetic protein-specific transcriptional activation of Id1 gene. J Biol Chem 2002 Feb 1; 277(5): 3176–3185.

8. Hjertner O, Hjorth-Hansen H, Börset M, Seidel C, Waage A, Sundan A. Bone morphogenetic protein-4 inhibits proliferation and induces apoptosis of multiple myeloma cells. Blood 2001 Jan 15; 97(2): 516–522.

9. Westhrin M, Holien T, Zahoor M, Moen SH, Buene G, Størdal B, et al. Bone Morphogenetic Protein 4 Gene Therapy in Mice Inhibits Myeloma Tumor Growth, But Has a Negative Impact on Bone. JBMR Plus 2020 Jan; 4(1): e10247.

10. Olsen OE, Wader KF, Misund K, Våtsveen TK, Rø TB, Mylin AK, et al. Bone morphogenetic protein-9 suppresses growth of myeloma cells by signaling through ALK2 but is inhibited by endoglin. Blood Cancer J 2014 Mar 21; 4(3): e196.

11. Holien T, Våtsveen TK, Hella H, Waage A, Sundan A. Addiction to c-MYC in multiple myeloma. Blood 2012 Sep 20; 120(12): 2450–2453.

12. Holien T, Våtsveen TK, Hella H, Rampa C, Brede G, Grøseth LA, et al. Bone morphogenetic proteins induce apoptosis in multiple myeloma cells by Smad-dependent repression of MYC. Leukemia 2012 May; 26(5): 1073–1080.

13. Bock C, Datlinger P, Chardon F, Coelho MA, Dong MB, Lawson KA, et al. High-content CRISPR screening. Nat Rev Methods Primers 2022; 2(1).

14. Iyer DN, Schimmer AD, Chang H. Applying CRISPR-Cas9 screens to dissect hematological malignancies. Blood Adv 2023 May 23; 7(10): 2252–2270.

15. Massari ME, Murre C. Helix-loop-helix proteins: regulators of transcription in eucaryotic organisms. Mol Cell Biol 2000 Jan; 20(2): 429–440.

16. Sun XH, Copeland NG, Jenkins NA, Baltimore D. Id proteins Id1 and Id2 selectively inhibit DNA binding by one class of helix-loop-helix proteins. Mol Cell Biol 1991 Nov; 11(11): 5603–5611.

17. Riechmann V, van Crüchten I, Sablitzky F. The expression pattern of Id4, a novel dominant negative helix-loop-helix protein, is distinct from Id1, Id2 and Id3. Nucleic Acids Res 1994 Mar 11; 22(5): 749–755.

18. Langlands K, Yin X, Anand G, Prochownik EV. Dimerential interactions of Id proteins with basic-helix-loop-helix transcription factors. J Biol Chem 1997 Aug 8; 272(32): 19785–19793.

19. Norton JD, Deed RW, Craggs G, Sablitzky F. Id helix-loop-helix proteins in cell growth and dimerentiation. Trends Cell Biol 1998 Feb; 8(2): 58–65.

20. Loveys DA, Streim MB, Kato GJ. E2A basic-helix-loop-helix transcription factors are negatively regulated by serum growth factors and by the Id3 protein. Nucleic Acids Res 1996 Jul 15; 24(14): 2813–2820.

21. Deed RW, Jasiok M, Norton JD. Lymphoid-specific expression of the Id3 gene in hematopoietic cells. Selective antagonism of E2A basic helix-loop-helix protein associated with Id3-induced dimerentiation of erythroleukemia cells. J Biol Chem 1998 Apr 3; 273(14): 8278–8286.

22. Burger R, Guenther A, Bakker F, Schmalzing M, Bernand S, Baum W, et al. Gp130 and ras mediated signaling in human plasma cell line INA-6: a cytokine-regulated tumor model for plasmacytoma. Hematol J 2001; 2(1): 42–53.

23. Behsen AD, Holien T, Micci F, Rye M, Rasmussen JM, Andersen K, et al. Cell surface marker heterogeneity in human myeloma cell lines for modeling of disease and therapy. Sci Rep 2024 Nov 20; 14(1): 28805.

24. Våtsveen TK, Børset M, Dikic A, Tian E, Micci F, Lid AH, et al. VOLIN and KJON-Two novel hyperdiploid myeloma cell lines. Genes Chromosomes Cancer 2016 Nov; 55(11): 890–901.

25. Roliński M, Montaldo NP, Aksu ME, Fordyce Martin SL, Brambilla A, Kunath N, et al. Loss of Mediator complex subunit 13 (MED13) promotes resistance to alkylation through cyclin D1 upregulation. Nucleic Acids Res 2021 Feb 22; 49(3): 1470–1484.

26. Li W, Xu H, Xiao T, Cong L, Love MI, Zhang F, et al. MAGeCK enables robust identification of essential genes from genome-scale CRISPR/Cas9 knockout screens. Genome Biol 2014; 15(12): 554.

27. Herrera B, Inman GJ. A rapid and sensitive bioassay for the simultaneous measurement of multiple bone morphogenetic proteins. Identification and quantification of BMP4, BMP6 and BMP9 in bovine and human serum. BMC Cell Biol 2009 Mar 19; 10: 20.

28. Burger R, Günther A, Klausz K, Staudinger M, Peipp M, Penas EM, et al. Due to interleukin-6 type cytokine redundancy only glycoprotein 130 receptor blockade emiciently inhibits myeloma growth. Haematologica 2017 Feb; 102(2): 381–390.

29. Olsen OE, Hella H, Elsaadi S, Jacobi C, Martinez-Hackert E, Holien T. Activins as Dual Specificity TGF-β Family Molecules: SMAD-Activation via Activin- and BMP-Type 1 Receptors. Biomolecules 2020 Mar 29; 10(4).

30. Olsen OE, Sankar M, Elsaadi S, Hella H, Buene G, Darvekar SR, et al. BMPR2 inhibits activin and BMP signaling via wild-type ALK2. J Cell Sci 2018 Jun 11; 131(11).

31. Quist-Løkken I, Andersson-Rusch C, Kastnes MH, Kolos JM, Jatzlau J, Hella H, et al. FKBP12 is a major regulator of ALK2 activity in multiple myeloma cells. Cell Commun Signal 2023 Jan 30; 21(1): 25.

32. Quist-Lokken I, Tilseth M, Andersson-Rusch C, Hanif MA, Walz M, Moen JFN, et al. Novel PROTAC to target FKBP12: the potential to enhance bone morphogenetic protein activity and apoptosis in multiple myeloma. Haematologica 2025 Apr 17.

33. Ruzinova MB, Benezra R. Id proteins in development, cell cycle and cancer. Trends Cell Biol 2003 Aug; 13(8): 410–418.

34. Kolos JM, Pomplun S, Jung S, Rieß B, Purder PL, Voll AM, et al. Picomolar FKBP inhibitors enabled by a single water-displacing methyl group in bicyclic [4.3.1] aza-amides. Chem Sci 2021 Nov 17; 12(44): 14758–14765.

35. Katagiri T, Imada M, Yanai T, Suda T, Takahashi N, Kamijo R. Identification of a BMP-responsive element in Id1, the gene for inhibition of myogenesis. Genes Cells 2002 Sep; 7(9): 949–960.

36. Kee BL, Rivera RR, Murre C. Id3 inhibits B lymphocyte progenitor growth and survival in response to TGF-beta. Nat Immunol 2001 Mar; 2(3): 242–247.

37. Jaleco AC, Stegmann AP, Heemskerk MH, Couwenberg F, Bakker AQ, Weijer K, et al. Genetic modification of human B-cell development: B-cell development is inhibited by the dominant negative helix loop helix factor Id3. Blood 1999 Oct 15; 94(8): 2637–2646.

38. Love C, Sun Z, Jima D, Li G, Zhang J, Miles R, et al. The genetic landscape of mutations in Burkitt lymphoma. Nat Genet 2012 Dec; 44(12): 1321–1325.

39. Ramis-Zaldivar JE, Gonzalez-Farré B, Balagué O, Celis V, Nadeu F, Salmerón-Villalobos J, et al. Distinct molecular profile of IRF4-rearranged large B-cell lymphoma. Blood 2020 Jan 23; 135(4): 274–286.

40. Li A, Yi H, Deng S, Ruan M, Xu P, Huo Y, et al. The genetic landscape of histologically transformed marginal zone lymphomas. Cancer 2024; 130(8): 1246–1256.

41. Lüscher B, Larsson LG. The basic region/helix-loop-helix/leucine zipper domain of Myc proto-oncoproteins: function and regulation. Oncogene 1999 May 13; 18(19): 2955–2966.

42. Roschger C, Cabrele C. The Id-protein family in developmental and cancer-associated pathways. Cell Commun Signal 2017 Jan 25; 15(1): 7.

43. Rowsey RA, Smoley SA, Williamson CM, Vasmatzis G, Smadbeck JB, Ning Y, et al. Characterization of TCF3 rearrangements in pediatric B-lymphoblastic leukemia/lymphoma by mate-pair sequencing (MPseq) identifies complex genomic rearrangements and a novel TCF3/TEF gene fusion. Blood Cancer J 2019 Oct 1; 9(10): 81.

44. Went M, Hoang PH, Law PJ, Kaiser MF, Houlston RS. Exploiting gene dependency to inform drug development for multiple myeloma. Sci Rep 2022 Jul 26; 12(1): 12696.

45. Kurata K, James-Bott A, Tye MA, Yamamoto L, Samur MK, Tai YT, et al. Prolyl-tRNA synthetase as a novel therapeutic target in multiple myeloma. Blood Cancer J 2023 Jan 12; 13(1): 12.

46. Agrawal P, Yu K, Salomon AR, Sedivy JM. Proteomic profiling of Myc-associated proteins. Cell Cycle 2010 Dec 15; 9(24): 4908–4921.

47. Gooding S, Olechnowicz SWZ, Morris EV, Armitage AE, Arezes J, Frost J, et al. Transcriptomic profiling of the myeloma bone-lining niche reveals BMP signalling inhibition to improve bone disease. Nat Commun 2019 Oct 4; 10(1): 4533.

