## Supplemental Table 1 for "Whole genome CRISPR knockout screen reveals ID3 as a key regulator of myeloma cell survival via TCF3 and c-MYC"

**Supplementary Table 1. Primers used in the study.**

Sequences of primers used in this study and their respective area of use.

| Name | Application | Sequence (5’ to 3’) |
| --- | --- | --- |
| 251_F1-Seq5N8 | 1st PCR | CTTTCCCTACACGACGCTCTTCCGATCTNNNNNNNNGTGGAA  AGGACGAAACACCG |
| 260_R1-Seq7N8 | 1st PCR | GGAGTTCAGACGTGTGCTCTTCCGATCTNNNNNNNNCGACT  CGGTGCCACTTTTTC |
| P1-A1 | 2nd PCR | CAAGCAGAAGACGGCATACGAGAT**GTCGGTAA**GTGACTGGA  GTTCAGACGTGTGCTCTTCCGATC*T |
| P2-A2 | 2nd PCR | CAAGCAGAAGACGGCATACGAGAT**AGGTCACT**GTGACTGGA  GTTCAGACGTGTGCTCTTCCGATC*T |
| P3-A3 | 2nd PCR | CAAGCAGAAGACGGCATACGAGAT**GAATCCGA**GTGACTGGA  GTTCAGACGTGTGCTCTTCCGATC*T |
| P4-A4 | 2nd PCR | CAAGCAGAAGACGGCATACGAGAT**GTACCTTG**GTGACTGGA  GTTCAGACGTGTGCTCTTCCGATC*T |
| P5-A5 | 2nd PCR | CAAGCAGAAGACGGCATACGAGAT**CATGAGGA**GTGACTGGA  GTTCAGACGTGTGCTCTTCCGATC*T |
| P6-A6 | 2nd PCR | CAAGCAGAAGACGGCATACGAGAT**TGACTGAC**GTGACTGGA  GTTCAGACGTGTGCTCTTCCGATC*T |
| P7-A7 | 2nd PCR | CAAGCAGAAGACGGCATACGAGAT**CGTATTCG**GTGACTGGA  GTTCAGACGTGTGCTCTTCCGATC*T |
| P8-A8 | 2nd PCR | CAAGCAGAAGACGGCATACGAGAT**CTCCTAGA**GTGACTGGA  GTTCAGACGTGTGCTCTTCCGATC*T |
| P9-A9 | 2nd PCR | CAAGCAGAAGACGGCATACGAGAT**TAGTTGCG**GTGACTGGA  GTTCAGACGTGTGCTCTTCCGATC*T |
| P10-A10 | 2nd PCR | CAAGCAGAAGACGGCATACGAGAT**GAGATACG**GTGACTGGA  GTTCAGACGTGTGCTCTTCCGATC*T |
| P11-A11 | 2nd PCR | CAAGCAGAAGACGGCATACGAGAT**AGGTGTAC**GTGACTGGA  GTTCAGACGTGTGCTCTTCCGATC*T |
| P12-A12 | 2nd PCR | CAAGCAGAAGACGGCATACGAGAT**TAATGCCG**GTGACTGGA  GTTCAGACGTGTGCTCTTCCGATC*T |
| P13-B1 | 2nd PCR | CAAGCAGAAGACGGCATACGAGAT**TCAGACGA**GTGACTGGA  GTTCAGACGTGTGCTCTTCCGATC*T |
| P14-B2 | 2nd PCR | 5CAAGCAGAAGACGGCATACGAGAT**GATAGGCT**GTGACTGGA  GTTCAGACGTGTGCTCTTCCGATC*T |
| P15-B3 | 2nd PCR | CAAGCAGAAGACGGCATACGAGAT**TGGTACAG**GTGACTGGA  GTTCAGACGTGTGCTCTTCCGATC*T |
| P16-B4 | 2nd PCR | CAAGCAGAAGACGGCATACGAGAT**CAAGGTCT**GTGACTGGA  GTTCAGACGTGTGCTCTTCCGATC*T |
| P17-B5 | 2nd PCR | CAAGCAGAAGACGGCATACGAGAT**GCTATCCT**GTGACTGGA  GTTCAGACGTGTGCTCTTCCGATC*T |
| P18-B6 | 2nd PCR | CAAGCAGAAGACGGCATACGAGAT**ATGGAAGG**GTGACTGGA  GTTCAGACGTGTGCTCTTCCGATC*T |
| P19-B7 | 2nd PCR | CAAGCAGAAGACGGCATACGAGAT**TCAAGGAC**GTGACTGGA  GTTCAGACGTGTGCTCTTCCGATC*T |
| P20-B8 | 2nd PCR | CAAGCAGAAGACGGCATACGAGAT**GTTACGCA**GTGACTGGA  GTTCAGACGTGTGCTCTTCCGATC*T |
| P21-B9 | 2nd PCR | CAAGCAGAAGACGGCATACGAGAT**AGTCTGTG**GTGACTGGA  GTTCAGACGTGTGCTCTTCCGATC*T |
| P31-C7 | 2nd PCR | CAAGCAGAAGACGGCATACGAGAT**AAGCACTG**GTGACTGGA  GTTCAGACGTGTGCTCTTCCGATC*T |
| P32-C8 | 2nd PCR | CAAGCAGAAGACGGCATACGAGAT**CTAGCAAG**GTGACTGGA  GTTCAGACGTGTGCTCTTCCGATC*T |
| P40-D4 | 2nd PCR | CAAGCAGAAGACGGCATACGAGAT**TCGGTTAC**GTGACTGGA  GTTCAGACGTGTGCTCTTCCGATC*T |
