## Supplemental Figures for "Whole genome CRISPR knockout screen reveals ID3 as a key regulator of myeloma cell survival via TCF3 and c-MYC"

**Supplementary Figures**

**Figure S1.** *ID3* and *ID1* mRNA expression in myeloma cell lines treated with BMP10.

**Figure S2.** Relative ID3 protein levels.

**Figure S3.** Significantly over- or underrepresented HLH proteins from the KO screen.

**Figure S4.** CRISPR gene dependencies.

**Supplementary Figure S1**

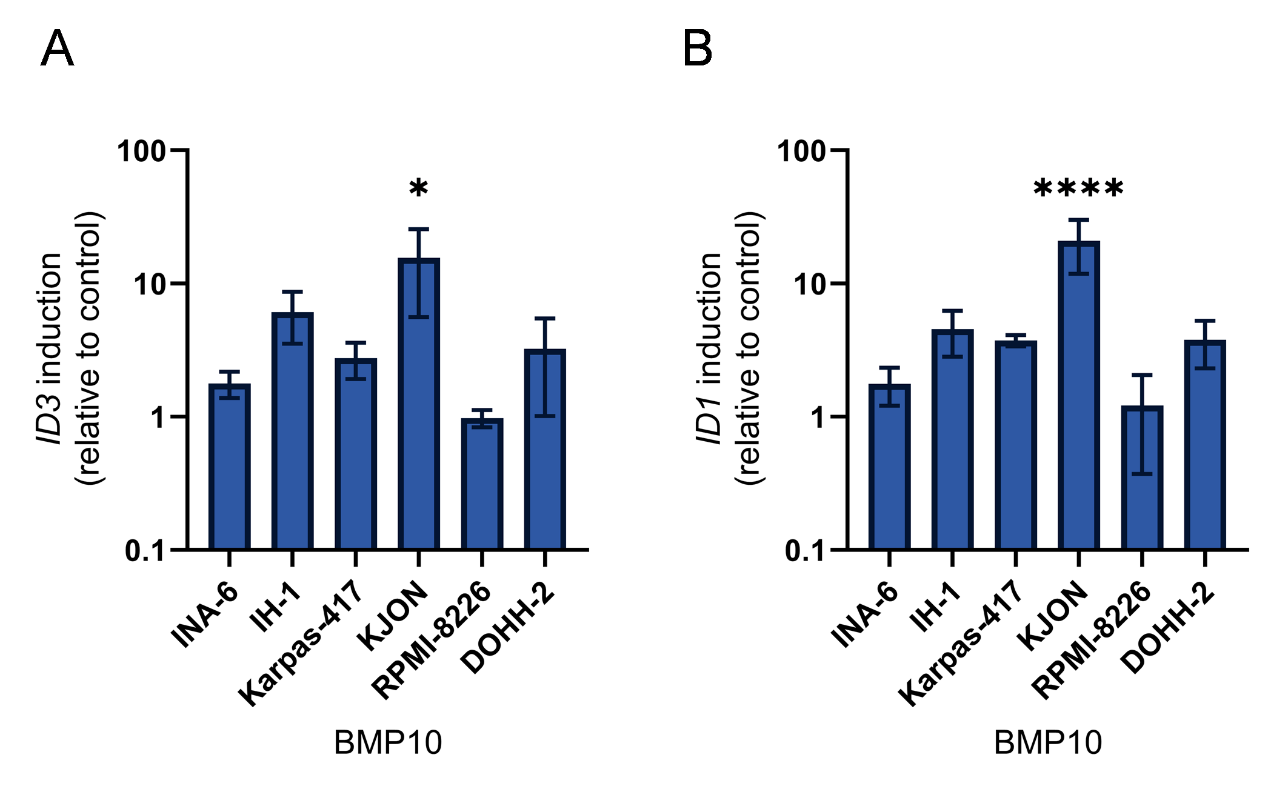

**Figure S1: ID3 and ID1 mRNA expression in myeloma cell lines treated with BMP10.** The cell lines INA-6, IH-1, Karpas-417, KJON, RPMI-8226, and DOHH-2 were treated with BMP10 (50 ng/mL) for 1 h and the mRNA levels of ID3 (A) and ID1 (B) were determined by RT-qPCR using n=3 independent experiments. The comparative Ct method was used with GAPDH as housekeeping gene.

**Supplementary Figure S2**

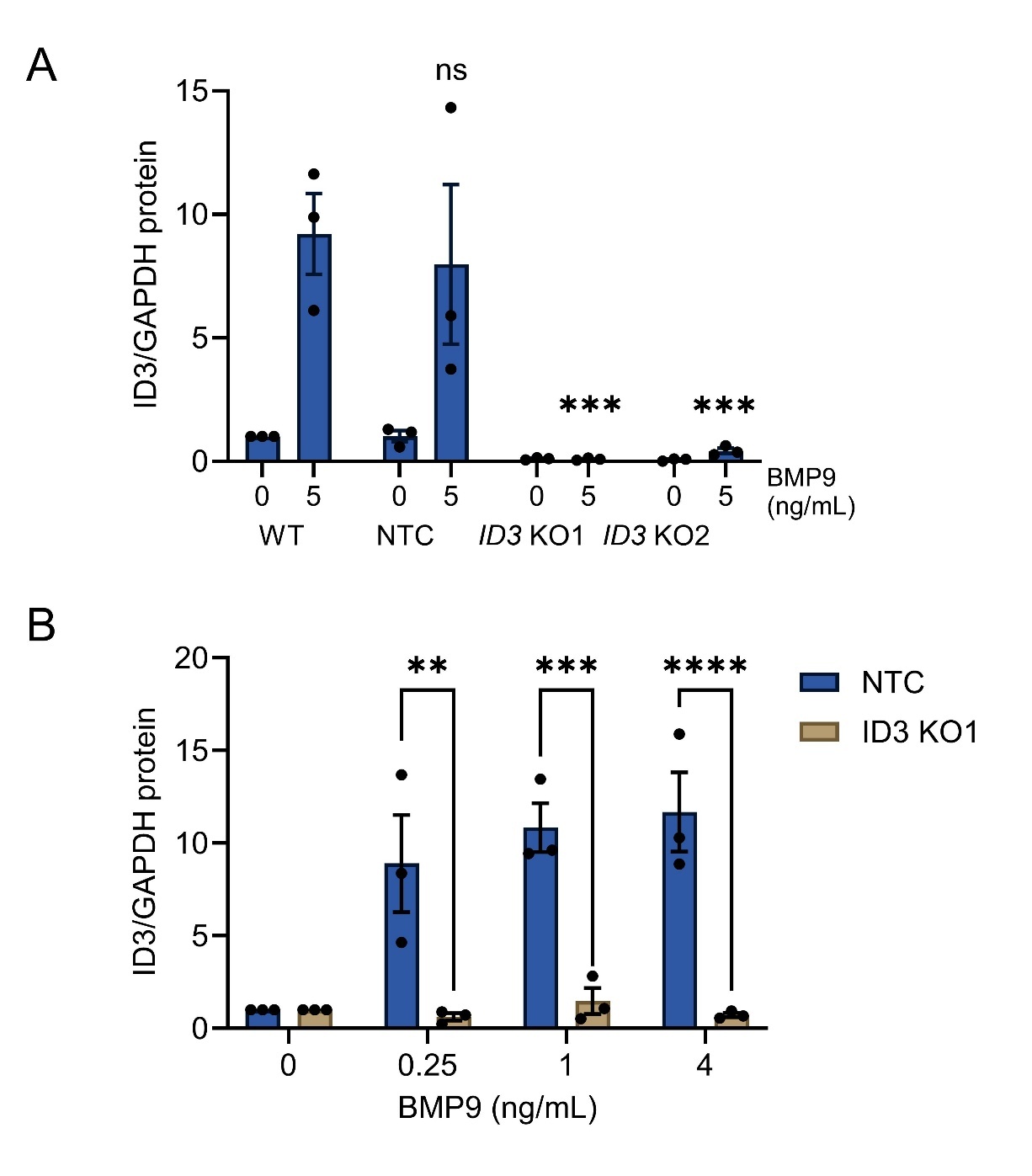

**Figure S2. Relative ID3 protein levels.** Densitometry of ID3 western blots in Fig. 3A (A) and Fig. 3F (B). The graphs show the combined results of n=3 independent experiments normalized to GAPDH and with WT (A) or NTC (B) cells. A compared the ID3 levels in BMP9-treated cells compared with WT as control, ***; p<0.001, two-way ANOVA, Dunnett’s multiple comparisons test and, B compared the relative induction of ID3 between NTC and ID3 KO1 cells, **; p<0.01, ***; p<0.001, ****; p<0.0001, two-way ANOVA, Šídák's multiple comparisons test.

**Supplementary Figure S3**

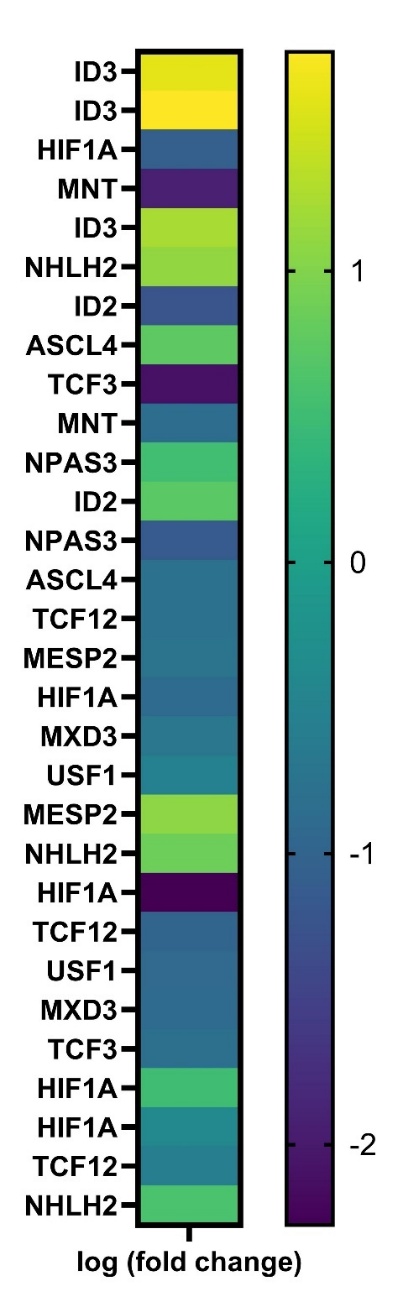

**Figure S3.** **Significantly over- or underrepresented HLH proteins from the KO screen.** The heatmap shows the genes encoding HLH proteins with significantly over- or underrepresented sgRNAs in the BMP9-treated versus control cells in the CRISPR/Cas9 KO screen. Each line represents one sgRNA and they are listed in the order of significance, with the most significant on top.

**Supplementary Figure S4**

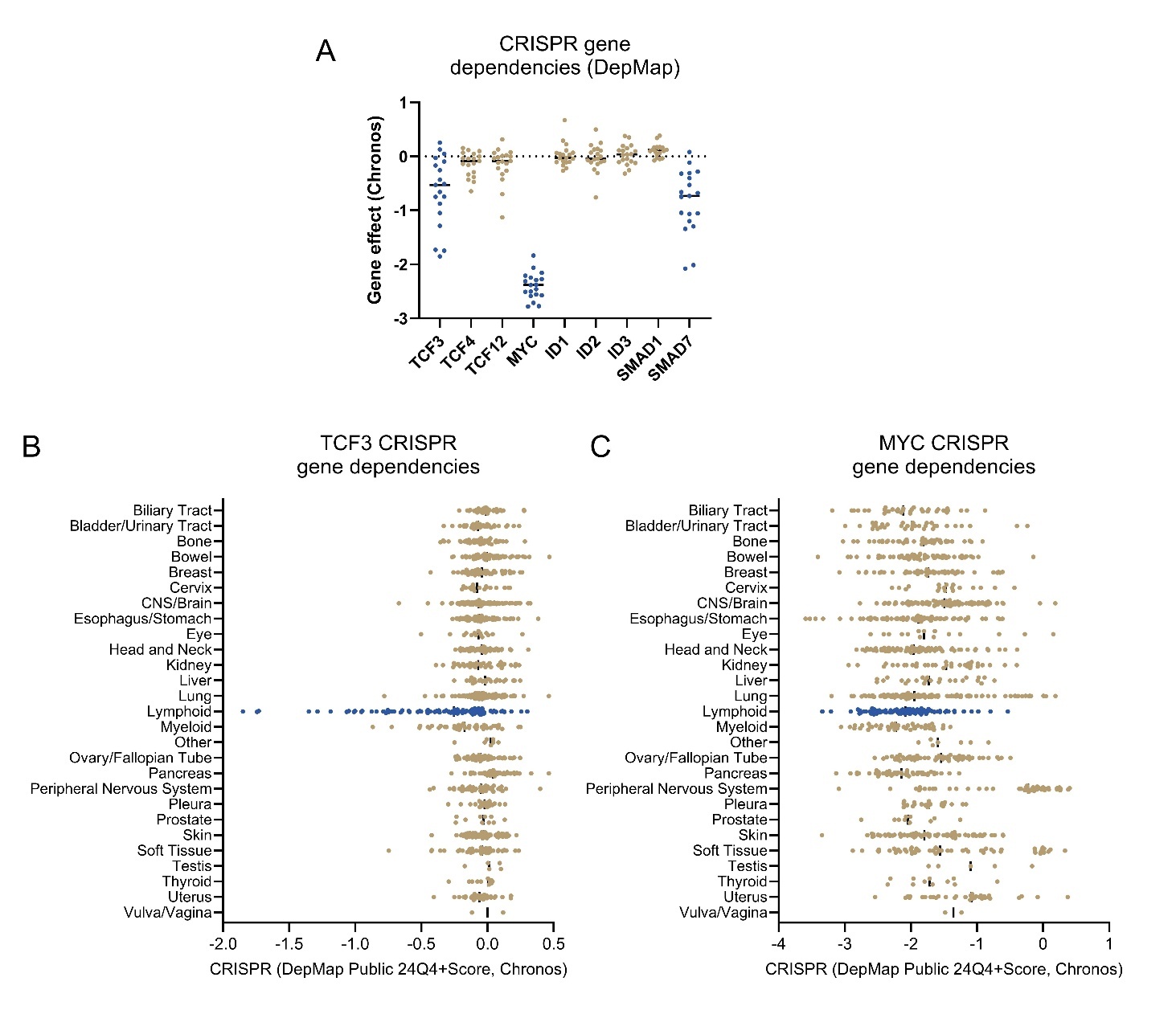

**Figure S4. CRISPR gene dependencies.** A. We used data from the DepMap Portal to compare dependencies of selected genes in the myeloma cell lines available in the database (n=19). We also looked at different cancer tissues and compared their dependencies for TCF3 (B) or MYC (C). TCF3 and MYC are highlighted in blue.
