## Supplemental Methods for "Whole genome CRISPR knockout screen reveals ID3 as a key regulator of myeloma cell survival via TCF3 and c-MYC"

**Supplementary Materials and methods**

**Cell growth conditions**

INA-6 was grown in 10 % heat-inactivated fetal calf serum (FCS) in RPMI-1640 (Sigma-Aldrich) with 2 mM L-glutamine (Sigma-Aldrich) (RPMI) and interleukin (IL)-6 (1 ng/mL) (Gibco, Thermo Fischer Scientific, Waltham, MA, USA). IH-1 cells were cultured using 10 % heat-inactivated human serum (HS) (Department of Immunology and Transfusion Medicine, St. Olav’s University hospital, Trondheim, Norway) in RPMI and IL-6 (2 ng/mL). KJON cells were grown in 5 % HS in RPMI with IL-6 (2 ng/mL). RPMI-8226 was grown in 20 % FCS in RPMI, and Karpas-417 and DOHH-2 were grown in 10 % FCS in RPMI. All cells were cultured at 37 °C in a humidified atmosphere containing 5 % CO2 and were regularly checked for mycoplasma.

**Apoptosis assay**

Flow cytometry was used to measure cell viability by labelling with Apotest Annexin A5-FITC kit (VPS Diagnostics, Hoeven, The Netherlands). In brief, cells were incubated with annexin V-FITC (0.2 µg/mL in binding buffer) for 1 h on ice. Propidium iodide (PI) (1.4 µg/mL) was added 5 min before running the samples using LSRII flow cytometer (BD Biosciences, San Jose, CA, USA).

**Cell viability and proliferation**

CellTiter-Glo (Promega, Madison, WI, USA) was used to measure ATP levels by luciferase in cells. If not otherwise indicated, 10 000 cells were seeded in 2 % HS in RPMI, with IL-6 (1 ng/mL) added for INA-6, in 96 well optical plates and treated as indicated. 70 µL CellTiter-Glo reagent was added to the cells, the plate was then shaken for 2 min and incubated for 10 min at room temperature. Plates were analyzed with Victor 1420 multilabel counter (PerkinElmer Inc. Waltham, MA, USA).

**INA-6 BRE-luc reporter assay**

The INA-6 BRE-luc reporter assay was performed as previously described.^1^

**Western blot**

Cells were treated as indicated, washed once with ice-cold PBS, and lysed for 30 min on ice. The lysis buffer consisted of 1 % IGEPAL-630 (Sigma-Aldrich), 150 mM NaCl, 50 mM Tris-HCl (pH 7.5), cOmplete Mini EDTA-free Protease Inhibitor Cocktail (#11836170001, Roche, Basel, Switzerland), 1 mM Na_3_VO_4_, and 50 mM NaF. For electrophoresis we used NuPAGE Bis-Tris gels and running buffer with MOPS or MES, depending on the protein sizes (Invitrogen, Thermo Fisher Scientific). Gels were blotted onto 0.45 µM nitrocellulose membranes, blocked with 5 % non-fat dry milk in Tris-buffered saline with 0.05 % Tween-20 (TBS-T). Primary antibodies used were phospho-SMAD1/5 (RRID: AB_491015, #9516), ID3 (RRID: AB_2732885, #9837), TCF3/E2A (RRID: AB_2797860, #12258), TCF12/HEB (RRID: AB_3101781, #95359), β-Actin (RRID: AB_2223172, #4970) all from Cell Signaling Technology (BioNordika AS, Oslo, Norway), c-Myc (RRID: AB_2148606, #551102)(BD Biosciences, BD Norge, Oslo, Norway), and glyceraldehyde-3-phosphate dehydrogenase (GAPDH) (RRID: AB_2107448, #Ab8245, Abcam, Cambridge, UK). The secondary antibodies used were either goat anti-rabbit or goat anti-mouse horseradish peroxidase conjugated (Dako Cytomation, Glostrup, Denmark). Positive bands were detected using the SuperSignal West Femto Maximum Sensitivity Substrate (#34096, Thermo Fisher Scientific) and ChemiDoc MP Imaging System (BioRad).

**RT-qPCR**

Cells were treated as indicated, washed once with ice-cold PBS before RNA was isolated with the RNeasy Mini Kit (#74106, Qiagen, Crawley, UK) using QIAcube automated sample preparation (Qiagen). cDNA synthesis was performed using the High-Capacity RNA-to-cDNA kit (#4388950, Applied Biosystems, Thermo Fisher Scientific). To perform RT-qPCR, we used standard settings of the StepOne Real-Time PCR system with PerfeCTa qPCR FastMix ROX (#733-1398, Quantabio, VWR, Oslo, Norway) and Taqman Gene Expression Assays, FAM-MGB labelled (Applied Biosystems, Thermo Fisher Scientific). Taqman assays used were *ID1* (Hs00357821_g1), *ID3* (Hs00171409_m1), *MYC* (Hs00153408_m1), and *GAPDH* (Hs99999905_m1). The comparative Ct method was used to calculate the relative changes in expression with GAPDH as the housekeeping gene.

**Lentiviral library generation**

HEK293T cells were seeded at 40 % confluence the day before transfection. Transfection was performed using VSV.G (a gift from Tannishtha Reya, Addgene plasmid #14888)^2^, psPAX2 (a gift from Didier Trono, Addgene plasmid #12260), and Human CRISPR Knockout Pooled Library (GeCKO v2) A and B (a gift from Feng Zhang (Addgene #1000000048)^3^, with Lipofectamine 2000 (#11668027, Invitrogen) according to the manufacturer’s protocol. The media was collected 59 h after transfection and viral particles were concentrated by ultracentrifugation (125 682 g, 2 h, 4 ℃). The lentiviral pellet was resuspended in DMEM with 1 % BSA (#15561020, Invitrogen), aliquoted and stored in -80 ℃ until use.

**Generation of single knockout cell lines**

For validation of the hits benchling.com was used to select two top-ranking sgRNAs not present in the GeCKO library, ID3 KO1 (CTGGTACCCGGAGTCCCGAG) and ID3 KO2 (GGTGCGCGGCTGCTACGAGG). As a negative control we used a non-targeting control sgRNA (NTC, CTTAAGTCATGAGCAAAGAT). Individual sgRNAs were cloned into plentiCRISPRv2 (a gift from Feng Zhang, Addgene plasmid #52961) following the standard protocol.^3^ Briefly, single oligos (Sigma Aldrich) with the 20 bp spacer sequence and overhangs were ligated. Double-stranded oligos were ligated into the BsmBI (Thermo Fisher Scientific) digested plentiCRISPRv2 backbone. Ligation mix was transformed into chemically competent E. coli DH5-α. Plasmid DNA was extracted with Midi kit (Macherey-Nagel) following the manufacturers protocol. Correct integration of the 20 bp spacer region was confirmed with PCR and gel electrophoresis. To generate knock-outs viral transduction with third generation lentiviral packaging plasmids using Genejuice (Novagen, Merck Life Science AS, Oslo, Norway) in 293T packaging cells (Open Biosystems, Thermo Fisher Scientific) was used. The supernatant was filtered and given to INA-6 cells in the presence of polybrene (8 µg/mL). Puromycin (0.5 µg/mL) was added 48 h after viral transduction to select for successful integration. Cells were single-cell sorted using BD FACSAria II Flow Cytometer (BD Biosciences) to a 96 well plate and expanded. Loss of protein expression was confirmed by western blotting.

**CoMMpass gene expression and overall survival analysis**

RNA sequencing expression data (E74GTF_Salmon_Gene_TPM) and overall survival (OS) data (STAND_ALONE_SURVIVAL) from the Multiple Myeloma Research Foundation (MMRF) CoMMpass IA16 release were downloaded from the MMRF webpage (http:://research.themmrf.org). For 767 patients, both RNA sequencing data from CD138+ cells at diagnosis and OS data were available. The patient samples were divided into above and below median TPM (transcript per million) expression of the specific gene of interest. Kaplan-Meier survival analyses were performed using GraphPad Prism version 9, and comparison of survival curves using the Log-rank Mantel-Cox test.

**Knockdown with siRNA**

For siRNA experiments, we transfected cells using the Nucleofector 2b device (Amaxa biosystems, Cologne, Germany) and Cell Line Nucleofector Kit R (Lonza, Basel, Switzerland) using a previously described protocol.^4^ The siRNAs used were ON-TARGETplus Non-targeting pool (D-001810-10-05) and ON-TARGETplus Human siRNAs for TCF3 (L-009384-00-0005), and TCF12 (L-006356-00-0005) (Dharmacon, Horizon Discovery, Cambridge, UK).

**Statistical analysis**

GraphPad Prism 10 (GraphPad Software, La Jolla, CA, USA) was used for calculations of statistical significance. The statistical tests used are described in the related methods’ paragraphs or in figure legends.

**References**

1.	Quist-Løkken I, Andersson-Rusch C, Kastnes MH, Kolos JM, Jatzlau J, Hella H, et al. FKBP12 is a major regulator of ALK2 activity in multiple myeloma cells. Cell Commun Signal 2023 Jan 30; 21(1): 25.

2.	Reya T, Duncan AW, Ailles L, Domen J, Scherer DC, Willert K, et al. A role for Wnt signalling in self-renewal of haematopoietic stem cells. Nature 2003 May 22; 423(6938): 409–414.

3.	Sanjana NE, Shalem O, Zhang F. Improved vectors and genome-wide libraries for CRISPR screening. Nat Methods 2014 Aug; 11(8): 783–784.

4.	Fagerli UM, Holt RU, Holien T, Vaatsveen TK, Zhan F, Egeberg KW, et al. Overexpression and involvement in migration by the metastasis-associated phosphatase PRL-3 in human myeloma cells. Blood 2008 Jan 15; 111(2): 806–815.
